# Genome-scale Capture C promoter interaction analysis implicates novel effector genes at GWAS loci for bone mineral density

**DOI:** 10.1101/405142

**Authors:** Alessandra Chesi, Yadav Wagley, Matthew E. Johnson, Elisabetta Manduchi, Chun Su, Sumei Lu, Michelle E. Leonard, Kenyaita M. Hodge, James A. Pippin, Kurt D. Hankenson, Andrew D. Wells, Struan F.A. Grant

**Affiliations:** Center for Spatial and Functional Genomics, Children’s Hospital of Philadelphia, Philadelphia, United States; Department of Orthopaedic Surgery, University of Michigan Medical School, Ann Arbor, United States; Institute for Biomedical Informatics, University of Pennsylvania Perelman School of Medicine, Philadelphia, United States; Department of Pathology and Laboratory Medicine, University of Pennsylvania Perelman School of Medicine, Philadelphia, United States; Department of Pediatrics, University of Pennsylvania Perelman School of Medicine, Philadelphia, United States; Divisions of Genetics and Endocrinology, Children’s Hospital of Philadelphia, Philadelphia, United States

## Abstract

Osteoporosis is a devastating disease with an essential genetic component. Genome wide association studies (GWAS) have discovered genetic variants robustly associated with bone mineral density (BMD), however they only report genomic signals and not necessarily the precise localization of culprit effector genes. Therefore, we sought to carry out physical and direct ‘variant to gene mapping’ in a relevant primary human cell type. We developed ‘SPATIaL-seq’ (genome-Scale, Promoter-focused Analysis of chromaTIn Looping), a massively parallel, high resolution Capture-C based method to simultaneously characterize the genome-wide interactions of all human promoters. By intersecting our SPATIaL-seq and ATAC-seq data from human mesenchymal progenitor cell -derived osteoblasts, we observed consistent contacts between candidate causal variants and putative target gene promoters in open chromatin for ~30% of the 110 BMD loci investigated. Knockdown of two novel implicated genes, *ING3* at ‘*CPED1-WNT16*’ and *EPDR1* at ‘*STARD3NL*’, had pronounced inhibitory effects on osteoblastogenesis. Our approach therefore aids target discovery in osteoporosis and can be applied to other common genetic diseases.

## INTRODUCTION

Osteoporosis is a common chronic form of disability due to loss of bone mineral density (BMD). During their lifetime, women lose 30–50% of peak bone mass, while men lose 20–30%. Fracture risk is higher in individuals with lower BMD^1-3^. Above age 50, many women of European ancestry will suffer at least one fracture; of these, many are at high risk for a subsequent fracture^4^. The subsequent loss in mobility and increased mortality have an enormous financial impact estimated at $17 billion annually^5^, and this is likely to rise during the next few decades due to an aging population^4^.

BMD is a classic complex trait influenced by behavioral, environmental and genetic factors. There is strong evidence for genetic predisposition to osteoporosis^6-8^, with an estimated 60% to 80% of the risk explained by heritable factors^9,10^. Population ancestry differences also speak to the genetic component^11,12^.

After the limited successes in the candidate gene^13,14^ and family-based linkage study eras of bone genetics^15,16^, the GWAS approach has proven a more comprehensive and unbiased strategy to identify loci related to this complex phenotype. Increasingly higher resolution GWAS have examined adult bone phenotypes^17-22^ - the latest meta-analysis reported 56 adult BMD (measured by dual-energy X-ray absorptiometry)^17-19^ and 14 fracture risk associated loci^22^, while additional loci have been discovered in the pediatric setting^23,24^. Indeed, we found that many of these adult bone loci also operate in childhood^23-27^, including rare variation at the engrailed 1 (*EN1*) locus^28^ uncovered in a recent sequencing study in a large sample of European adults^29^.

However, GWAS only reports the sentinel single nucleotide polymorphism (SNP), i.e. the SNP at a given locus with the lowest association *P*-value, which is unlikely to be the actual causal variant. Furthermore, the locations of the GWAS signals, the vast majority of which are non-coding in nature, residing in either intronic or intergenic regions, do not necessarily imply the precise location of the underlying effector genes. Thus, key questions related to GWAS are: is the nearest gene to a GWAS-implicated SNP in fact the actual principal culprit gene at the locus? Or is it another gene somewhere in the neighborhood? One example of this is the characterization of the key *FTO* locus in obesity^30-32^. The top GWAS signal resides within an intronic region of the *FTO* gene but in fact is primarily driving the expression of the *IRX3* and *IRX5* genes nearby, i.e. this variant appears to be in an embedded enhancer in one gene but influencing the expression of others. As a second example, we implicated a nearby gene, *ACSL5*^33^, at the key type 2 diabetes *TCF7L2* locus^34,35^, with GTEx confirming a co-localized eQTL for this gene^36^. This should not be too surprising given that gene expression can be controlled locally or via long range interactions over large genomic distances. After all, it is well established that in non-coding regions of the genome there are important regulatory elements, such as enhancers and silencers, and genetic variants that disrupt those elements can equally confer susceptibility to complex disease. Indeed, many regulatory elements do not control the nearest genes and can reside tens or hundreds of kilobases away.

Given this, there is a compelling need to systematically characterize the mechanisms of action at each of the BMD GWAS loci, in order to identify the effector genes. Because of the paucity of public domain genomic data relevant to bone, such as eQTL and chromatin conformation capture data, we elected to evaluate key BMD GWAS signals^22-24,27,37,38^ in the context of next level, three-dimensional genomics by leveraging a Capture C-based high resolution promoter ‘interactome’ methodology (SPATIaL-seq). Chromatin conformation capture-based techniques, combined with ATAC-seq, can help to establish initial ‘variant-to-gene mapping’, from which therapeutic and diagnostic approaches can be developed with greater confidence given that the correct target is actually being pursued. With the need for functional insight into GWAS observations, our goal was to provide the first comprehensive ‘variant to gene mapping’ for BMD GWAS-implicated loci by leveraging human MSC-derived osteoblasts, a relevant cellular model for human bone biology.

## RESULTS

In order to investigate the genome-wide contacts of all promoters in the human genome (i.e. a promoter ‘interactome’), we developed a new capture-C-based approach (‘SPATIaL-seq’) based on a custom-designed SureSelect library (Agilent) targeting the transcriptional start sites (TSS) of 22,647 coding genes and 21,684 non-coding RNAs (including their alternative promoters). We adapted existing Capture-C protocols^39,40^ using a 4-cutter restriction enzyme (DpnII, average fragment size 433 bp) to achieve higher resolution than the more commonly used 6-cutter (HindIII, average fragment size 3.7 kb). The design included 36,691 bait fragments associated to 44,331 transcripts, representing the most comprehensive 3D promoter interactome analyzed to date.

The premise to utilizing a combination of SPATIaL-seq and ATAC-seq was to get as close to the BMD GWAS causal variant as possible through wet lab determination in a cell line relevant for human bone biology, as opposed to statistical ascertainment - there is currently a paucity of genotyped trans-ethnic cohorts with BMD data to conduct fine mapping. We therefore carried out comprehensive ATAC-seq and SPATIaL-seq in order to physically fine-map each BMD locus in primary human MSC-derived osteoblasts.

Primary MSCs were cultivated using standard techniques and then osteoblast induction was performed with BMP2, as previously described^41^. ATAC-seq data from nine libraries derived from four donors (**Table S1**) were analyzed using the ENCODE ATAC-seq pipeline (https://github.com/kundajelab/atac_dnase_pipelines) yielding 156,406 open chromatin “conservative” peaks, which allowed determination of informative (i.e. residing in open chromatin) proxy SNPs for each of the 110 independent (r^2^<0.2) signals at 107 BMD GWAS loci (**Table S2**). This was accomplished by overlapping the positions of the open chromatin regions (peaks) with those of the sentinel and proxy SNPs (r^2^>0.4 to sentinel SNP in Europeans; total n=14,007 proxies) at each of the BMD GWAS loci. This effort substantially advanced the initial GWAS discoveries, since ATAC-seq permitted us to identify a shortlist of 474 candidate variants residing within open chromatin that are in LD with the sentinel SNP of 88 out of the 107 BMD loci investigated.

Next, we performed SPATIaL-seq on MSC-derived osteoblasts from three donors (**Table S1**). Libraries from each donor yielded high coverage (an average of ~1.6 billion reads per library) and good quality, with >40% valid read pairs and >75% capture efficiency (% of unique valid reads captured; T**able S3**). We called significant interactions using the CHiCAGO pipeline^42^. We first performed analyses at 1-fragment and 4-fragment resolutions and observed that the median distance for *cis* interactions increased when decreasing the resolution (**Table S4**). Therefore, we merged the results from the two analyses, allowing us to use a very high resolution for short-distance interactions, and to trade off resolution for increased sensitivity at longer interaction distances. Using this approach, we identified a total of 295,422 interactions (~14% were bait to bait), with a median distance for *cis* interactions of 50.5 kb, and a low number of trans interactions (0.7%). Most of the non-bait promoter-interacting regions (PIRs) had contacts with a single baited region (84%), while only 1% contacted more than four.

PIRs were significantly enriched for open chromatin regions as detected by our ATAC-seq experiments, suggesting a potential regulatory role. They were also enriched for histone marks associated with active chromatin regions in primary human osteoblasts from the ENCODE project^43^ (**Table S5**), such as enhancer regions (H3K27ac and H3K4me1), active promoters (H3K4me3), actively transcribed regions (H3K36me3), and transcription factor binding sites (H3K4me2); they also associated with CTCF binding sites and the repressive mark H3K27me3, but not with the repressive mark H3K9me3 (**Fig S1A)**. PIRs were highly enriched for BMD GWAS signals and their proxies (r^2^>0.4), but substantially less so for the non bone-related Alzheimer’s disease (**Fig S1B**). The number of contacts per bait were also significantly higher (*P*<2×10^-16^) in open vs inaccessible promoters, as determined by ATAC-seq, but only when considering contacts to ‘open’ PIRs.

To explore the relationship between promoter interactivity measured by SPATIaL-seq and gene expression, we performed RNA-seq on MSC-derived osteoblasts from the same three donors. Markers of osteoblast differentiation such as *SPP1, SOST, DKK1, Osterix* (*SP7*), and *DMP1*, were abundantly expressed (>50^th^ percentile) in all three samples (highlighted in red in **Table S6**). We observed an increased number of contacts in the more highly expressed genes, but only when considering open promoters and open PIRs, as determined by our ATAC-seq experiments (**Fig. S2**).

Of the BMD GWAS loci with an open chromatin SNP in MSC-derived osteoblasts (not residing in a baited promoter region), 33 revealed a significant direct interaction to an ‘open’ promoter region **(Table 1**).

**Table 1.**
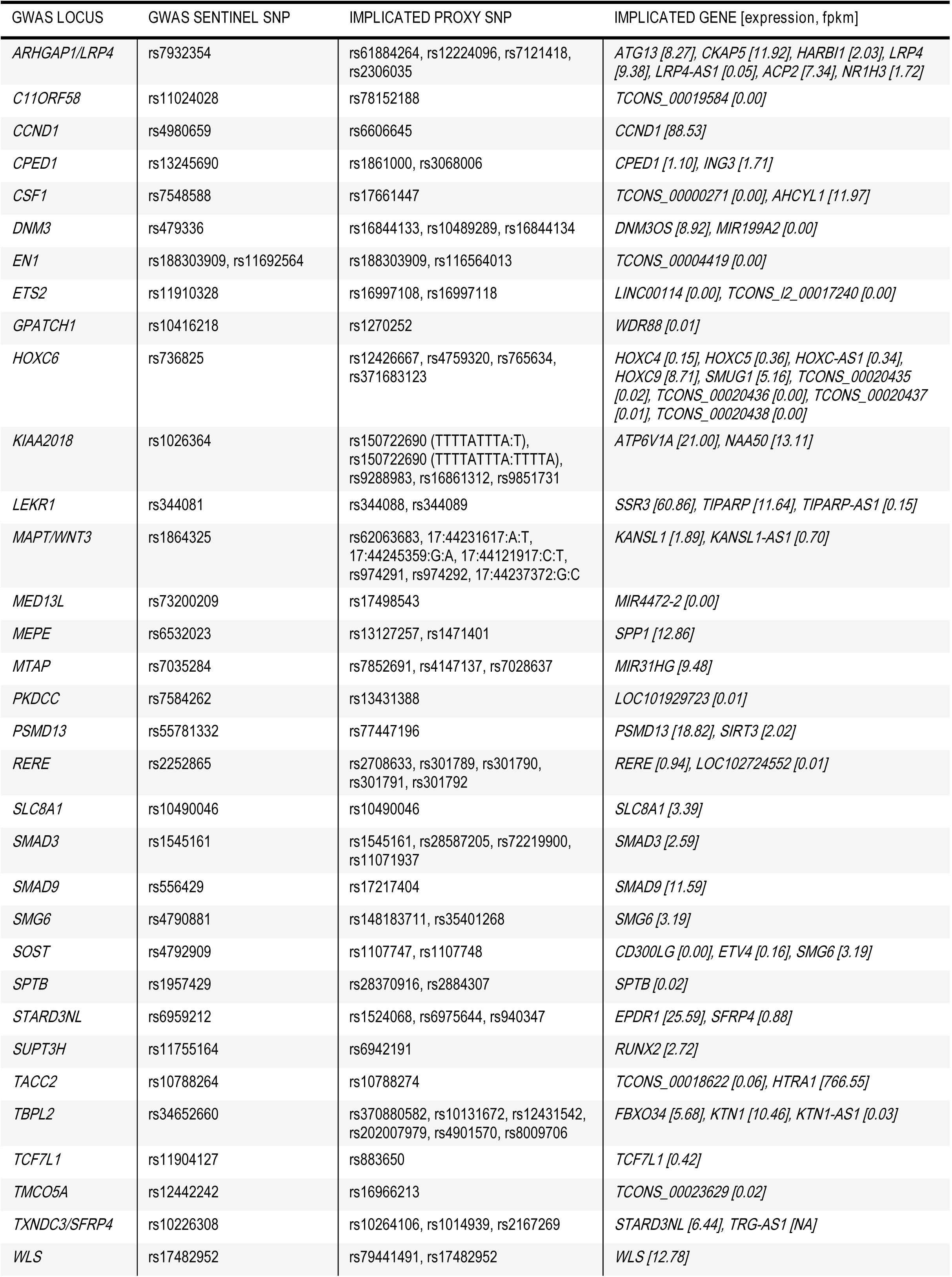
Implicated SNPs and target genes at 33 BMD GWAS loci in hMSC-derived osteoblasts. For each locus, proxy SNPs in open chromatin looping to an open promoter are reported, together with their sentinel SNP and the associated GWAS locus. Looping interactions and open chromatin maps were derived from SPATIaL-seq and ATAC-seq experiments on hMSC-derived osteoblasts cultures from 3 or more individuals. Average expression (in fpkm) of the target genes in hMSC-derived osteoblasts from RNA-seq experiments from the same 3 individuals is also reported.

We detected a total of 80 distinct chromatin looping interactions, involving 60 baited regions corresponding to 65 ‘open’ promoters and 66 open chromatin regions harboring one or more BMD proxy SNPs (22 interactions were detected at both1-fragment and 4-fragment resolution, 6 at 1-fragment resolution only, and 52 at 4-fragment only; **Table S7**). The vast majority of the implicated genes were highly expressed in the MSC-derived osteoblasts (**Table S6**).

SNP-promoter interactions (for SNPs not residing in baits) fell in to three types of observations: 1. to nearest gene only (36%) (**Fig. 1A**), to both nearest and more distant gene(s) (12%) (**Fig. 1B**) and only to distant gene(s) (52%) (**Fig. 1C**). Overall, 58% of the GWAS loci interacted with only one baited promoter region, while 42% interacted with more than one.

**Figure 1.**
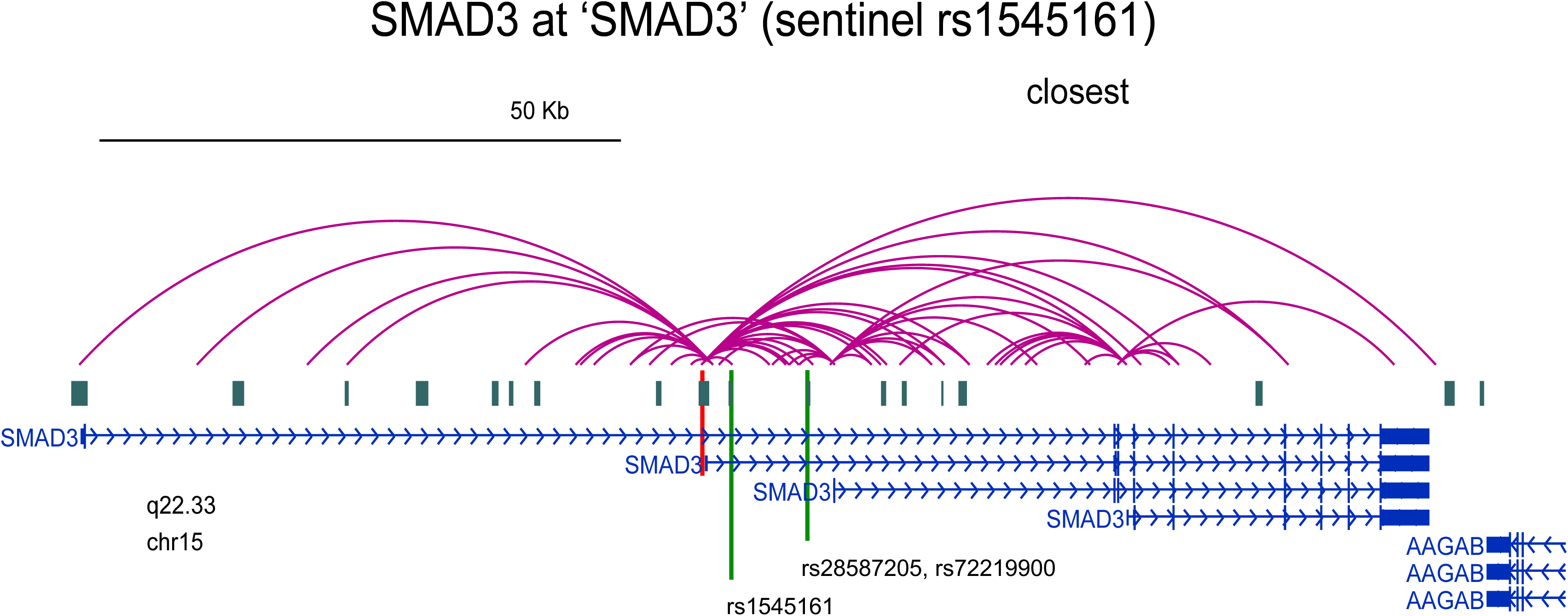
Examples of interactions between ATAC-seq implicated BMD SNPs and open promoters detected by SPATIaL-seq. **(A)** to nearest gene only. **(B)** to both nearest and more distant gene(s). **(C)** only to distant gene(s). Contacts were visualized using the WashU EpiGenome Browser. Red lines indicate genes of interest, while the green lines represent the sentinel and key proxy SNPs.

We went on to carry out pathway analyses on the above genes implicated by ATAC-seq plus SPATIaL-seq in human MSC-derived osteoblasts. Using Ingenuity, we found four enriched canonical pathways surviving multiple comparison correction, all very relevant to osteoblastic differentiation: ‘osteoarthritis pathway’, ‘TGF-β signaling’, ‘role of osteoblasts, osteoclasts and chondrocytes in rheumatoid arthritis’ and ‘BMP signaling pathway’ (**Fig. S3**). Among the implicated genes, we found several with a known role in osteogenesis, such as *RUNX2* at the ‘*SUPT3H*-*RUNX2’* locus, *HTRA1*^*44*^ at ‘*TACC2*’ and *MIR31HG*^*45*^ at ‘*MTAP*’, confirming the validity of our approach, while other target genes were completely novel.

In order to validate our findings, we targeted the expression of implicated genes using siRNA at two key loci in primary human MSCs derived from four donors (**Table S1**): *WNT16, CPED1* and *ING3* at the ‘*WNT16-CPED1*’ locus, and *EPDR1* and *SFRP4* at the ‘*STARD3NL*’ locus, and then assessed osteoblast differentiation. qPCR analysis showed that each siRNA resulted in significant knockdown of its corresponding target across the donor MSCs, but did not impact the expression of the other gene or genes implicated at the same loci (**Fig. 2 B, C; Fig. 3 B, C, D**). BMP2 treatment had no effect on *ING3* expression, but reduced *CPED1* and increased *WNT16*. In addition, upon BMP2 treatment, basal expression of *SFRP4* decreased, while *EPDR1* increased, although, considered across donors (which show variability), the results are not statistically significant. Since ALP expression is considered essential for hydroxyapatite deposition and hard-tissue mineralization, histochemical ALP staining was performed using parallel sets of gene-targeted cells. While targeting *WNT16, CPED1* and *SFRP4* produced somewhat variable ALP staining across donor lines, staining was strikingly and consistently reduced by *ING3* and *EPDR1* targeting. Furthermore, Alizarin red S staining confirmed that calcium phosphate mineral deposition was absent from cells with decreased *ING3* and *EPDR1* expression, whereas it could be observed in cells that lacked *CPED1, WNT16* and *SFRP4* expression. These changes in osteoblast differentiation while associated with reduced ALP gene expression (**Fig. 2G and 3H**), were not associated with concomitant decreases in osteoblast transcriptional regulator Osterix nor did the knockdowns negatively impact BMP2 signaling based on Id1 expression. Surprisingly, despite the negative impact of *ING3* knockdown on osteoblast differentiation, *ING3* knockdown increased both Id1 expression and Runx2 expression more than control siRNA. Runx2 expression was not significantly affected by *EPDR1* knockdown (although expression was variable among the samples).

**Figure 2.**
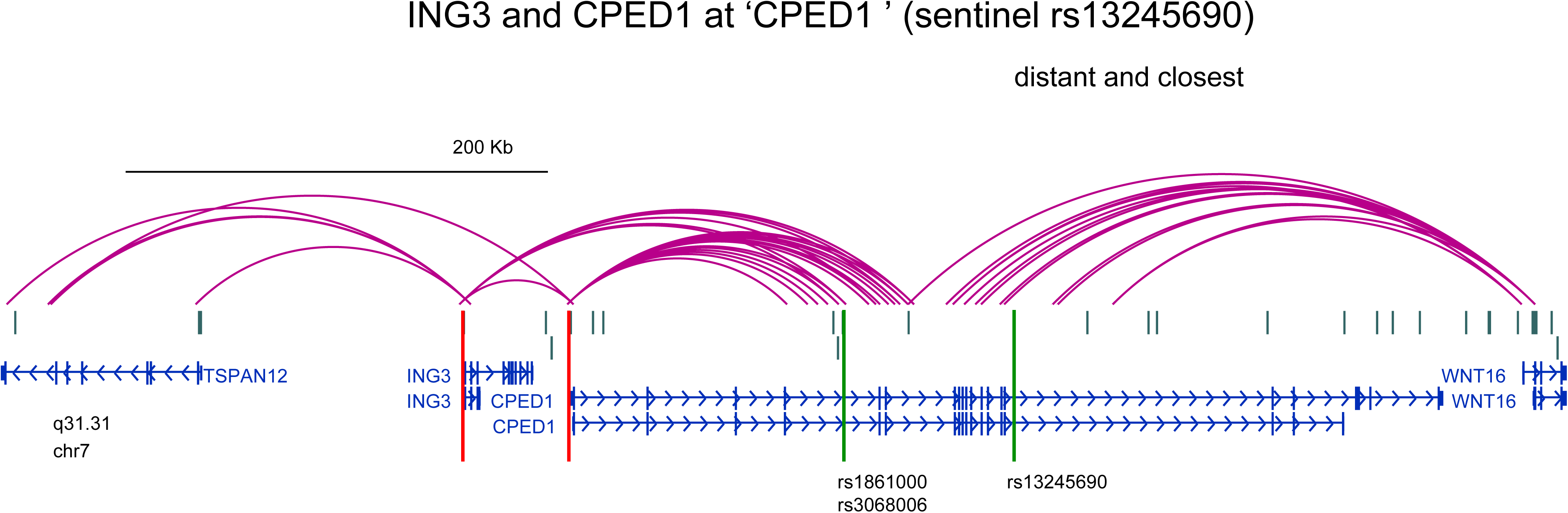
Knockdown of *ING3* gene expression - but not for *WNT16* or *CPED1* - impairs osteoblast differentiation. *CPED1* is implicated as an osteoporosis-associated gene by GWAS, but SPATIaL-seq data suggests that *ING3* may be a more likely causative gene. *ING3* knockdown results in a complete disruption of alkaline phosphatase induction and Alizarin red S staining, but there is no effect of *CPED1* and *WNT16* knockdown. (A) Representative AlkPhos (purple) and Alizarin (red) stained plates (repeated with 4 different independent hMSC donor cell lines). (B-H) Quantitative gene expression. Grey columns = No BMP treatment; Black columns = BMP treatment. Columns = mean. Error bars = Standard deviation. n=3-4 unique donor lines. *, p<0.05 comparing No treatment to BMP treatment for each siRNA. #, p<0.05 comparing control siRNA to siRNA for gene of interest.

**Figure 3.**
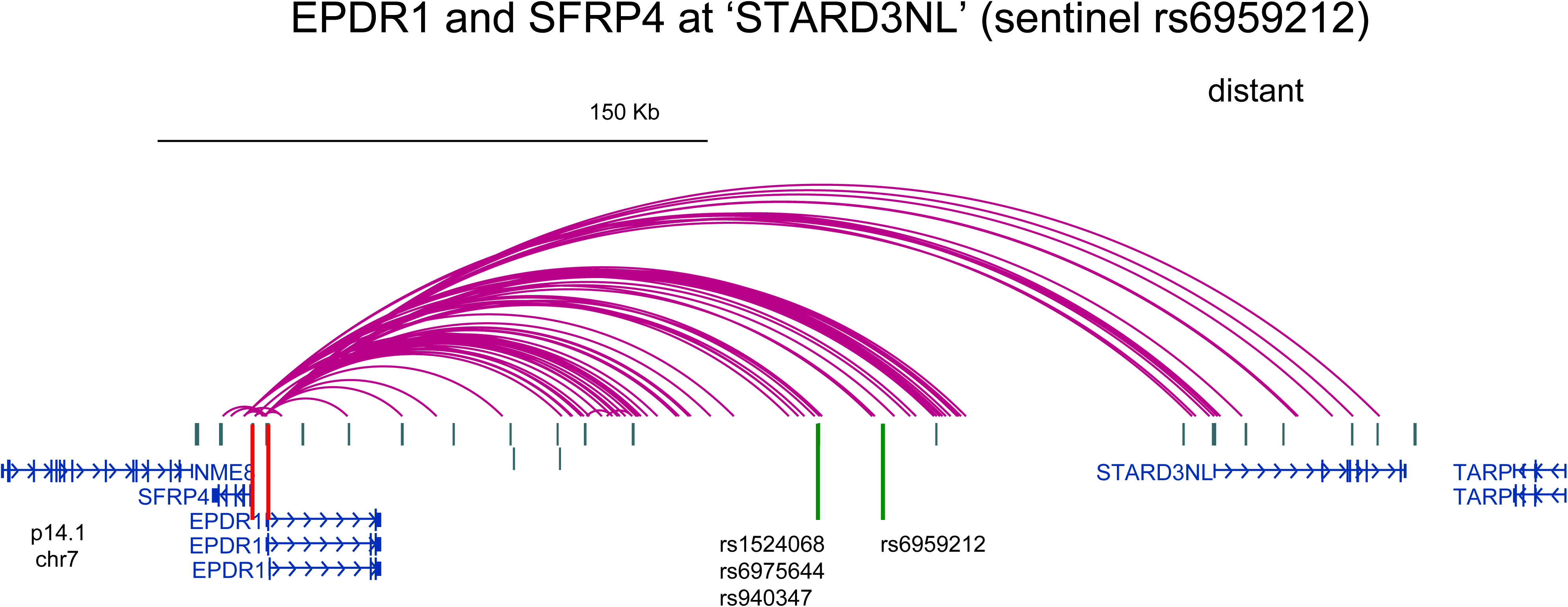
Knockdown of *EPDR1* gene expression - but not for *SFRP4* - impairs osteoblast differentiation. *EPDR1* disruption results in a reduction in alkaline phosphatase expression and activity and alizarin red S staining, but there is no effect of *SFRP4* knockdown. (A) Representative AlkPhos (ALP) (purple) and Alizarin (red) stained plates (repeated with 4 different independent hMSC donor cell lines). (B-G) Quantitative gene expression. Grey columns = No BMP treatment; black columns = BMP treatment. Columns = mean. Error bars = Standard deviation. n=3-4 unique donor lines. *, p<0.05 comparing No treatment to BMP treatment for each siRNA. #, p<0.05 comparing control siRNA to siRNA for gene of interest.

In summary, knock-down of two novel genes (*ING3* and *EPDR1*) not previously associated with BMD but implicated by our combined ATAC-seq and SPATiAL-seq approach revealed strong effects on osteoblast differentiation (decreased ALP expression and absence of calcium phosphate mineral deposition), suggesting an important role in bone biology and possibly in the development of diagnostic and therapeutic tools for osteoporosis.

## DISCUSSION

Recently, many chromatin conformation capture (3C) based methods have been developed with the goal of mapping GWAS variants to their target genes^39,40,46-49^. While Hi-C was developed to study the high order genomic organization of human chromatin domains, and not the precise looping interactions between GWAS-implicated variants and their target genes (and requires a high amount of sequencing), the main limitations of the other currently available approaches are the low resolution (dictated by the use of the 6-cutter HindIII) of Capture Hi-C^39^, and the bias intrinsic to the Hi-ChIP technique^49^, which is not ‘promoter-centric’ and uses antibodies to capture genomic regions characterized by certain histone marks (such as H3K27ac). In addition, none of these methods has been designed to target promoters of non-coding genes or alternative promoters for the same gene.

To overcome these limitations, and given the paucity of bone-related cell types represented in the public domain (ENCODE, GTEx^36^ etc.), we developed SPATIaL-seq, a very nimble high-resolution technique that does not require the large sample sizes typically required for eQTL analysis, providing significant results with only a few biological replicates. Moreover, SPATIaL-seq allows for a non-hypothesis driven gene discovery effort, since it presumes nothing about the region in which the SNP resides.

In this study, we applied SPATIaL-seq to primary human MSC-derived osteoblasts, representing a highly relevant, but relatively difficult to obtain, cellular model type to study BMD. We recognize that other cell types, most particularly osteoclasts, are also involved in BMD determination, but we have shed light on how these loci operate in this key single cell type, most responsible for building peak bone density.

We were able to determine complex and intricate contacts for ~30% of the BMD loci, frequently detecting two or more promoter contacts at these signals. About 80% of the ATAC-seq implicated BMD proxy SNPs not residing in a promoter interacted with distal genes, either exclusively or in conjunction with an interaction with the nearest gene. Implicated genes revealed enrichment for functionally relevant networks for bone biology. We went on to demonstrate with siRNA mediated knockdown experiments that *ING3* and *EPDR1*, implicated genes at the ‘*WNT16-CPED1*’ and ‘*STARD3NL*’ loci, respectively, play a role in osteoblast differentiation. Most intriguingly, neither gene is widely known to the bone community and thus reveals novel biology.

Although our studies have not probed the mechanistic basis for the regulation of osteoblastogenesis by *ING3* and *EPDR1*, prior studies in other organ systems may provide some clues regarding potential function. Two mechanisms can be proposed by which *ING3* affects osteoblast differentiation. First, since the ING3 protein is part of the NuA4 histone acetyltransferase (HAT) complex that recognizes trimethylated forms of lysine 4 of histone H3 (H3K4me3)^50^, silencing of *ING3* might affect key osteoblastic genes during human osteoblast differentiation. Second, given that ING3 levels are down-regulated in head and neck carcinoma, melanoma, ameloblastoma, hepatocellular carcinoma, and colorectal cancers^50^, and ING3 containing transcription complexes interact with p^53^ transactivated promoters including promoters of p21/waf1 and Bax to affect cell cycle progression^51^, the absence of ING3 may allow cells to avoid differentiation by continuously re-entering the cell cycle; however, our studies are carried out in serum-free media and thus, we did not note any changes in cell proliferation. EPDR1 is a type II transmembrane protein that is similar to two families of cell adhesion molecules, the protocadherins and ependymins that can affect the extracellular milieu (ECM). Calcium-induced conformational change of EPDR1 molecules are known to be important for EPDR1 interaction with the components of the extracellular matrix and this can affect cell adhesion and migration^52^. Notably, extracellular matrix components play an important role in osteoblast differentiation, and thus, alterations in interactions of EPDR1 with the ECM could have important implications for this biological process. Intriguingly, an *Epdr1* KO mouse exists in the Mouse Genome Database (MGD) Project and has a short tibia phenotype.

We cannot exclude the role of other genes at these loci. At ‘*WNT16-CPED1*’, the culprit gene is usually believed to be *WNT16*, since the WNT pathway plays a fundamental role in bone development and Wnt16 has been shown to regulate cortical bone mass and bone strength in mice^53^. Indeed, two SNPs in LD with the *WNT16* GWAS sentinel rs3801387 (rs142005327, r^2^=0.94; rs2908004, r^2^=0.45) reside in the *WNT16* promoter region and may well regulate WNT16 expression; however, our SPATIaL-seq data reveal no looping between this promoter and putative enhancer open chromatin regions in our cell setting.

At the ‘*STARD3NL*’ locus, a compelling candidate effector gene is *SFRP4*, encoding secreted frizzled-related protein 4, a soluble Wnt inhibitor recently implicated in bone remodeling both in mouse and humans^54,55^. Nonetheless, and most interestingly, our approach was able to uncover the role of two less obvious, novel genes in osteoblastogenesis and bone mineralization.

In conclusion, we observed consistent contacts at multiple BMD GWAS loci with a high resolution promoter interactome applied to a single difficult to obtain, disease-relevant cell type i.e. human primary MSC-derived osteoblasts. Our ATAC-seq plus SPATIaL-seq approach has promise for future efforts to implicate effector genes at GWAS loci for other common genetic disorders.

## METHODS

### Loci analyzed

We leveraged 63 loci from the latest GWAS of adult BMD and fracture^27^, plus pediatric and adult loci from studies from our and other investigators’ groups^23,24,27,29,38,56,57^, for a total of 110 independent sentinels (**Table S2**). To obtain proxy SNPs, we used rAggr (http://raggr.usc.edu) with an r^2^<0.4 threshold and the all European (CEU+FIN+GBR+IBS+TSI) population. For two loci (*IZUMO* and *DHH*) for which we could not find proxies in rAggr, we leveraged the WTCCC genotype data (ref) to calculate r^2^ in PLINK v 1.9 (http://pngu.mgh.harvard.edu/purcell/plink/; ref), with a 0.4 threshold. For one low MAF locus (*LDLRAD3*) we could not identify proxies by either approach, so we ultimately worked with 14,007 proxies for 110 sentinels at 107 loci.

### Culturing, differentiation and functional characterization of hMSCs

Primary bone-marrow derived human MSCs isolated from healthy donors (age range: 22 years-29 years) were characterized for cell surface expression (CD166+CD90+CD105+/CD36-CD34-CD10-CD11b-CD45-) and tri-lineage differentiation (osteoblastic, adipogenic and chondrogenic) potential at the Institute of Regenerative Medicine, Texas A&M University. Expansion and maintenance of the cells were carried out using alpha-MEM supplemented with 16.5% FBS in standard culture conditions by plating cells at a density of 3000 cells/cm^2^. For osteoblastic differentiation, 15,000 cells/cm^2^ from maintenance cultures were plated in alpha-MEM consisting of 16.5% FBS, 25 μg/ml Ascorbic acid-2-phosphate, 5 mM beta-glycerophosphate and 1% insulin-transferrin-selenous acid (osteogenic media) and stimulated the next day with recombinant human BMP2 (300 ng/ml) (R&D Systems, MN) in serum-free osteogenic media. Cells were harvested at 72 hours following BMP2 treatment for sequencing library preparations because our previous work has shown that this time point reflects a stage when the cells are fully osteoblast committed but have not begun to mineralize. It was important to harvest non-mineralizing cultures, given that during that terminal state cells will begin to undergo apoptosis ^41,58^. Cells were assessed for differentiation in parallel plates by harvesting RNA and assessing the expression of RUNX2, SP7, and alkaline phosphatase, and target genes. Additionally, a third parallel plate of cells was assessed for alkaline phosphatase activity. For quantitative RT-PCR analysis, 300 ng of purified total RNA was reverse transcribed using High Capacity cDNA Reverse Transcription Kit (Applied Biosystems) in a 20 μl reaction. 1 μl of the resulting cDNA was amplified using Power SYBR^®^ Green PCR Master Mix and gene-specific primers in a 7500 Fast Real-Time PCR System (Applied Biosystems) following manufacturer’s recommendations. For assessing mineralization, cells were analyzed 8-10 days after BMP stimulation by staining with Alizarin red S. All values are reported as mean ± standard deviation with statistical significance determined via 2-way homoscedastic Student’s t-tests (*P≤0.05, #P≤0.10, N.S. = “not significant”).

### ATAC-seq library generation and peak calls

Following Tn5 Transposes transposition (Illumina Cat #FC-121-1030, Nextera) and purification of Tn5 Transposes derived DNA library in Michigan of native chromatin in 100,000 human MSC-derived osteoblasts for efficient epigenomic profiling of open chromatin, the samples were shipped to the Center of Spatial and Functional Genomics at CHOP where we completed the ATAC-seq process. This allowed for systematic insight in to which of the SNPs under investigation exhibit open chromatin. ATAC-seq is known to be compatible with many methods for cell collection and works effectively for many cell types. The methodology begins with harvesting live cells via trypsinization, followed by a series of wash steps. 100,000 cells of each were spun down at 550 ×g for 5 min, 4°C. The cell pellet was then resuspended in 50 μl cold lysis buffer (10 mM Tris-HCl, pH 7.4, 10 mM NaCl, 3 mM MgCl2, 0.1% IGEPAL CA-630). We spun down immediately at 550 ×g for 10 min, 4°C and then resuspended the nuclei in the transposition reaction mix (2x TD Buffer (Illumina Cat #FC-121-1030, Nextera), 2.5ul Tn5 Transposes (Illumina Cat #FC-121-1030, Nextera) and Nuclease Free H_2_O) on ice then incubated the transposition reaction at 37°C for 45 min. The transposed DNA was then purified using a Qiagen MinElute Kit with 10.5 μl elution buffer, frozen and sent to the Center for Spatial and Functional Genomics at CHOP. The transposed library was then PCR amplified using Nextera primers for 12 cycles. The PCR reaction was subsequently cleaned up using AMPureXP beads (Agencourt) and then paired-end sequenced on an Illumina HiSeq 4000 (100 bp read length) and the Illumina NovaSeq platform. Open chromatin regions were then called using the ENCODE ATAC-seq pipeline (https://www.encodeproject.org/atac-seq/) and selecting the resulting IDR conservative peaks. We define a genomic region ‘open’ if it has 1 bp overlap with an ATAC-seq peak.

### Cell fixation for chromatin capture

The protocol used for cell fixation was similar to previous methods^40^. Cells were collected and single-cell suspension were made with aliquots of 10^7^ cells in 10ml media (e.g. RPMI + 10%FCS). 540 μl 37% formaldehyde was added and incubation was carried out for 10 min at RT in a tumbler. The reaction was quenched by adding 1.5ml 1M cold glycine (4°C) giving a total 12ml. Fixed cells were centrifuged for 5 min at 1000 rpm at 4°C, and supernatant were removed. The pellets were washed in 10 ml cold PBS (4°C) by centrifugation for 5 min at 1000 rpm at 4°C. Supernatant was removed and cell pellets were resuspended in 5 ml of cold lysis buffer (10 mM Tris pH8, 10 mM NaCl, 0.2% NP-40 (Igepal) supplemented with protease inhibitor cocktails). Resuspended cells were incubated for 20 minutes on ice and centrifuged to remove the lysis buffer. Finally, the pellets were resuspended in 1 ml lysis buffer and transferred to 1.5ml Eppendorf tubes prior to snap freezing (ethanol / dry ice or liquid nitrogen). Cells were stored at -80 °C at this point until they were thawed again for digestion.

### 3C library generation

Initial 3C libraries were generated for fixed MSC-derived osteoblasts shipped from Michigan to CHOP. For each library, 10 million cells were harvested and fixed. The DNA was digested using DpnII, then re-ligated together using T4 DNA ligase and finally isolated by phenol/chloroform extraction^40^. In line with the previously published Capture C protocol^40^ the above described 3C libraries were utilized for the capture procedure.

The protocol used for 3C library generation was similar to previous methods^40^. Cell were thawed on ice, spun down and the lysis buffer was removed. The pellet was resuspended in water and incubated on ice for 10 minutes, followed by centrifugation and removal of supernatant. The pellet was then resuspended with 20% SDS and 1X NEBuffer DpnII and incubated at 37 °C for 1 h at 1,000 r.p.m. on a MultiTherm (Sigma-Aldrich). It was then further incubated for another 1 hour after the addition of Triton X-100 (concentration, 20%). After the 1 hour incubation 10 μL 50 U/μL DpnII (NEB) was add and left to digest until the end of the day. An additional 10 μL DpnII was added and digestion was left overnight at 37 °C. The next day, another 10 μL of DpnII was added and incubated for a further 3 hours.

The chromatin was then ligated overnight (8 μL T4 DNA Ligase, HC ThermoFisher (30 U/μL); with final concentration, 10 U ml and shaken at 16 °C at 1,000 r.p.m. on the MultiTherm. The next day, an additional 2 μL T4 DNA ligase was spiked in to each sample, and incubated for 3 more hours. The ligated samples were then decrosslinked overnight at 65 °C with Proteinase K (Invitrogen) and the following morning incubated for 30 min at 37 °C with RNase A (Millipore). Phenol-chloroform extraction was then performed, followed by an ethanol precipitation overnight at -20°C and then washed with 70% ethanol. Digestion efficiencies of 3C libraries were assessed by gel electrophoresis on a 0.9% agarose gel and quantitative PCR (SYBR green, Thermo Fisher).

### SPATIaL-seq

Custom capture baits were designed using Agilent SureSelect library design targeting both ends of DpnII restriction fragments encompassing promoters (including alternative promoters) of all human coding genes, noncoding RNA, antisense RNA, snRNA, miRNA, snoRNA, and lincRNA transcripts, totaling 36,691 RNA baited fragments. The capture library design covered 95% of all coding RNA promoters and 88% of RNA types describe above. The missing 5% of coding genes that weren’t able to be designed were either duplicated genes or contained highly repetitive DNA in their promoter regions.

The isolated DNA of the 3C libraries generated by DpnII digestion and ligation were quantified using a Qubit fluorometer (Life technologies), and 10μg of each library was sheared in dH2O using a QSonica Q800R to an average DNA fragment size of 350bp. QSonica settings used were 60% amplitude, 30 seconds on, 30 seconds off, 2 minute intervals, for a total of (5 intervals) at °C. After shearing, DNA was purified using AMPureXP beads (Agencourt), the concentration was checked via Qubit and DNA size was assessed on a Bioanalyzer 2100 using a 1000 DNA Chip. Agilent SureSelect XT Library Prep Kit (Agilent) was used to repair DNA ends and for adaptor ligation following the standard protocol. Excess adaptors were removed using AMPureXP beads. Size and concentration are checked again before hybridization. 1ug of ligated library was used as input for the SureSelect XT capture kit using their standard protocol and our custom-designed Capture-C library. The quality of the captured library was assessed by Bioanalyser 2100Qubit and Bioanalyzer using a high sensitivity DNA Chip. Each SureSelect XT library was initially sequenced on 1 lane HiSeq 4000 paired-end sequencing (100 bp read length) for QC. All 6 Capture C promoter ‘interactome’ libraries were then sequenced three at a time on an S2 flow cell on an Illumina NovaSeq, generating ~1.6 billion paired-end reads per sample.

### Analysis of SPATIaL-seq data

Quality control of the raw fastq files was performed with FastQC. Paired-end reads were pre-processed with the HICUP pipeline^59^, with bowtie2 as aligner and hg19 as reference genome. Significant promoter interactions at 1-DpnII fragment resolution were called using CHiCAGO^42^ with default parameters except for binsize which was set to 2500. Signifcant interactions at 4-DpnII fragment resolution were also called with CHiCAGO using artificial .baitmap and .rmap files where DpnII fragments were grouped into 4 consecutively and using default parameters except for removeAdjacent which was set to False. Results from the two resolutions were merged by taking the union of the interaction calls at either resolution and removing any 4-fragment interaction which contained a 1-fragment interaction. We define PIR a promoter-interacting region, irrespective of whether it is a baited region or not. The CHiCAGO function peakEnrichment4Features() was used to assess enrichment of genomic features in promoter interacting regions at both 1-fragment and 4-fragment resolution. Finally, we made use of the Washington Epigenome Browser (https://epigenomegateway.wustl.edu) to visualize the detected interactions within the context of other relevant functional genomics annotations.

### RNA-seq

Total RNA was isolated from differentiating osteoblasts using TRIzol reagent (Invitrogen) following manufacturer instructions, and then depleted of rRNA utilizing the Ribo-Zero rRNA Removal Kit (Illumina). RNA-seq libraries were prepared using the NEBNext Ultra II Directional RNA Library Prep Kit for Illumina (NEB) following standard protocols. Libraries were sequenced on one S2 flow cell on an Illumina NovaSeq 6000, generating ~200 million paired-end 50bp reads per sample. RNA-seq data were aligned to the hg19 genome with STAR v. 2.5.2b^60^ and pre-processed with PORT (https://github.com/itmat/Normalization) using the GENCODE Release 19 (GRCh37.p13) annotation plus annotation for lincRNAs and sno/miRNAs from the UCSC Table Browser (downloaded 7/7/2016). Normalized PORT counts for the uniquely mapped read pairs to the sense strand were additionally normalized by gene size and the resulting values were used in the computation of gene expression percentiles.

### Experimental knockdown of candidate genes and functional characterization

We investigated five key loci - *EPDR1* and SFRP4 at the ‘*STARD3NL*’ locus and WNT16, CPED1 and ING3 at the ‘*WNT16-CPED1*’ locus. Experimental knock down of these genes was achieved using DharmaFECT 1 transfection reagent (Dharmacon Inc., Lafayette, CO) using a set of 4 ON-TARGETplus siRNA (target sequences in **Table S1**) in three temporally separated independent MSC-derived osteoblast samples and then assessed for metabolic and osteoblastic activity. Following siRNA transfection, cells were allowed to recover for 2 days and stimulated with BMP2 for additional 3 days in serum-free osteogenic media, as previously described, after which the influence of knockdown on gene expression (qPCR) and early osteoblast differentiation (ALP) were evaluated as described.

## SUPPLEMENTAL CAPTIONS

**Figure S1.**
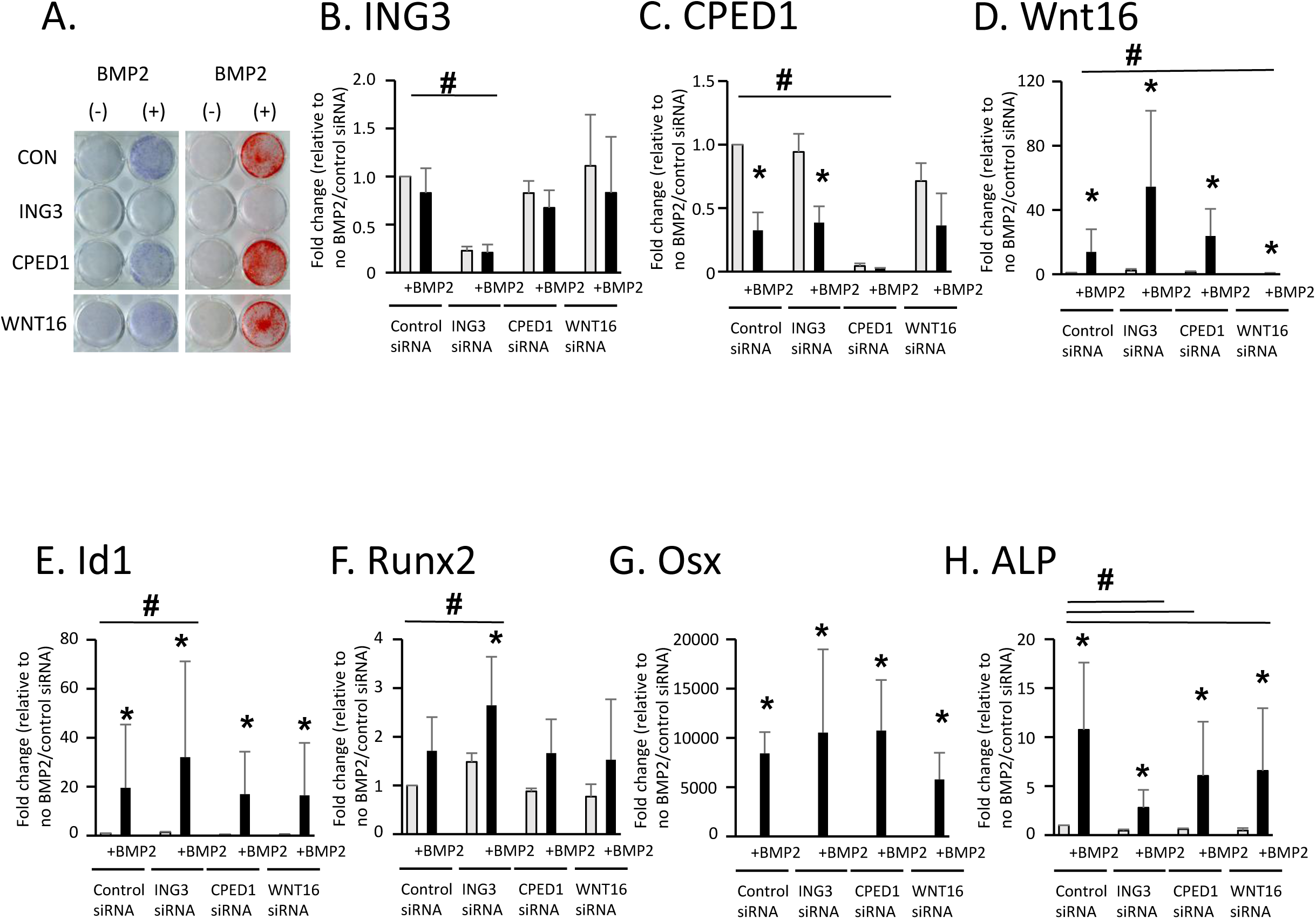
(A) Enrichment of promoter-interacting regions (PIR) in various chromatin marks. Yellow bars: number of overlaps with significant (CHiCAGO score >5) cis-interacting fragments (at 4-fragment resolution; bait-to-bait interactions were excluded); blue bars: expected overlaps based on 100 random subsets of fragments with a similar distribution of distances from the baits. ATAC-seq data is from our own experiments in hMSC-derived osteoblasts; other markers are from primary human osteoblasts from the ENCODE project. Error bars represent 95% confidence intervals. **(B) Enrichment of promoter-interacting fragments in BMD GWAS signals.** As above, but overlaps are computed with BMD or Alzheimer’s disease GWAS SNPs and their proxies (r^2^>0.4).

**Figure S2.**
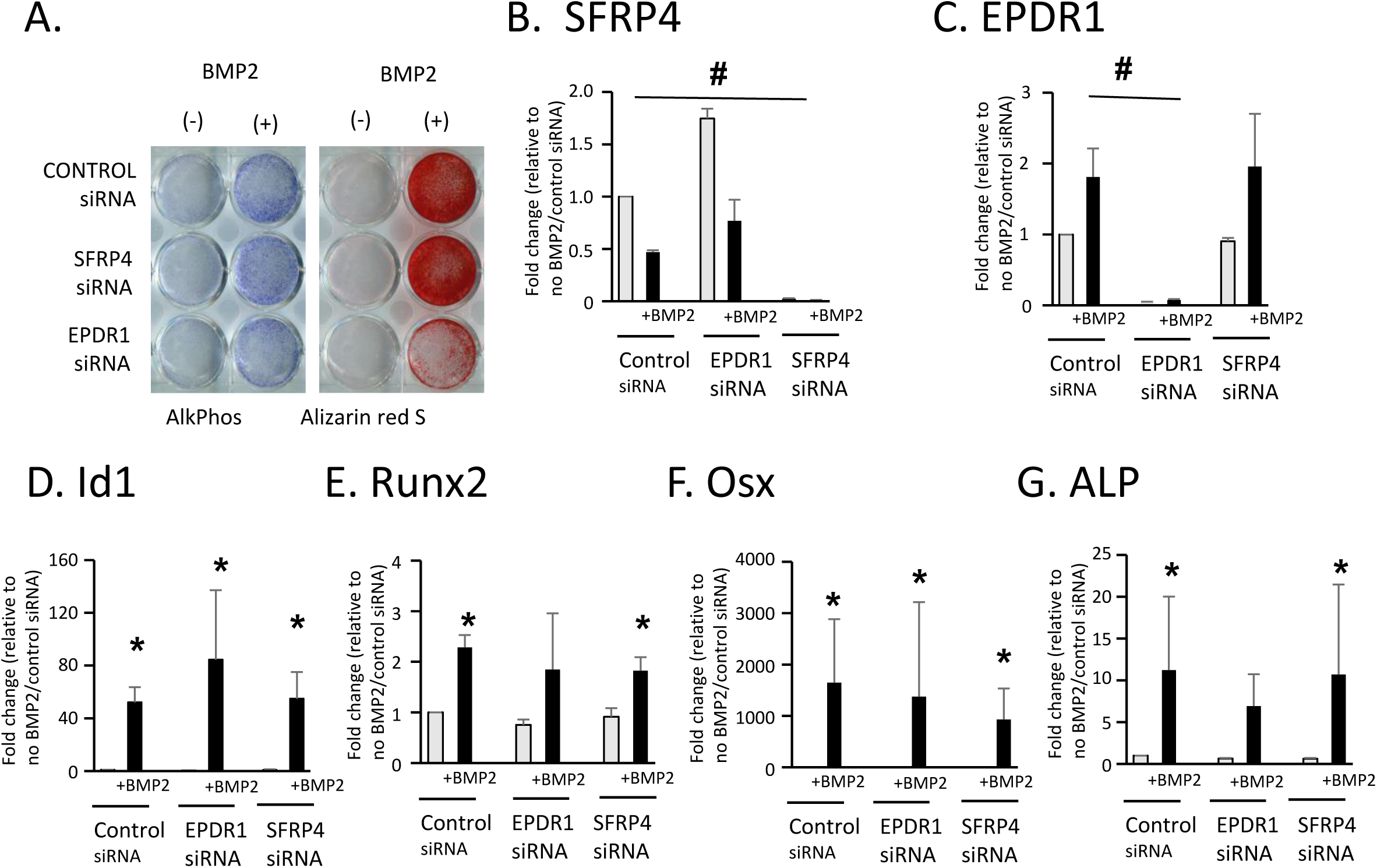
The number of contacts per bait correlates with gene expression, but only when considering open promoters and open PIR. Boxplots of the number of (cis) contacts per bait (at 4-fragment resolution; bait-to-bait interactions were excluded) for each quintile of expression of the baited gene(s). The horizontal line represents the median, the lower and upper hinges correspond to the 25th and 75th percentiles, and outliers (> 1.5 * IQR from the hinges, indicated by the whiskers) are plotted as single dots. **(A)** All baits and PIR. **(B)** Open baits and open PIR only.

**Figure S3.**
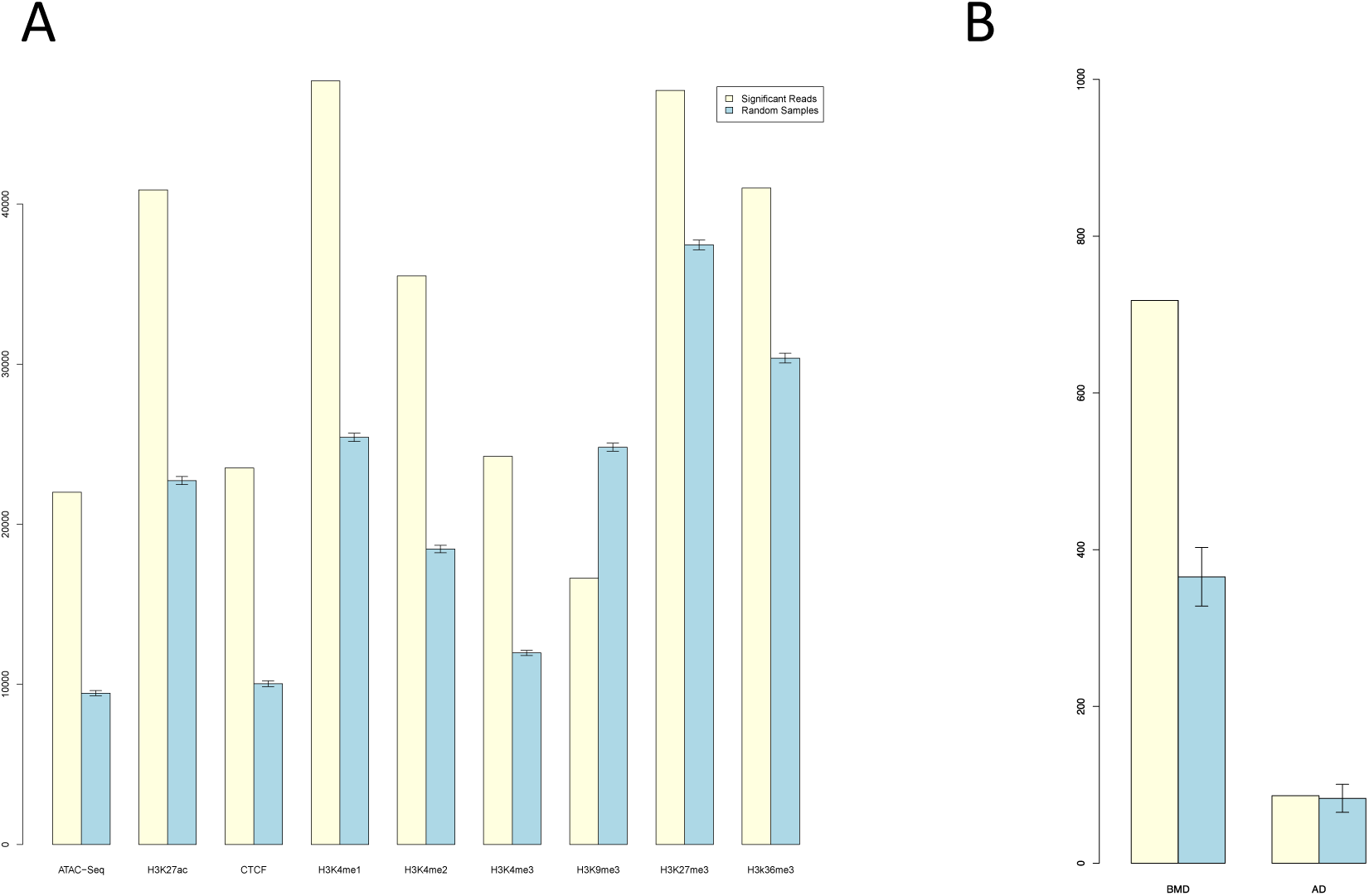
Canonical pathways enriched in the genes implicated by ATAC-seq plus SPATIaL-seq in human MSC-derived osteoblasts. The network shows each pathway as a single “node” colored proportionally to the B-H multiple testing corrected p-value, where brighter red = more significant. A line connects any two pathways when there are at least two implicated genes in common between them. The analysis was performed using Ingenuity Pathway Analysis software.

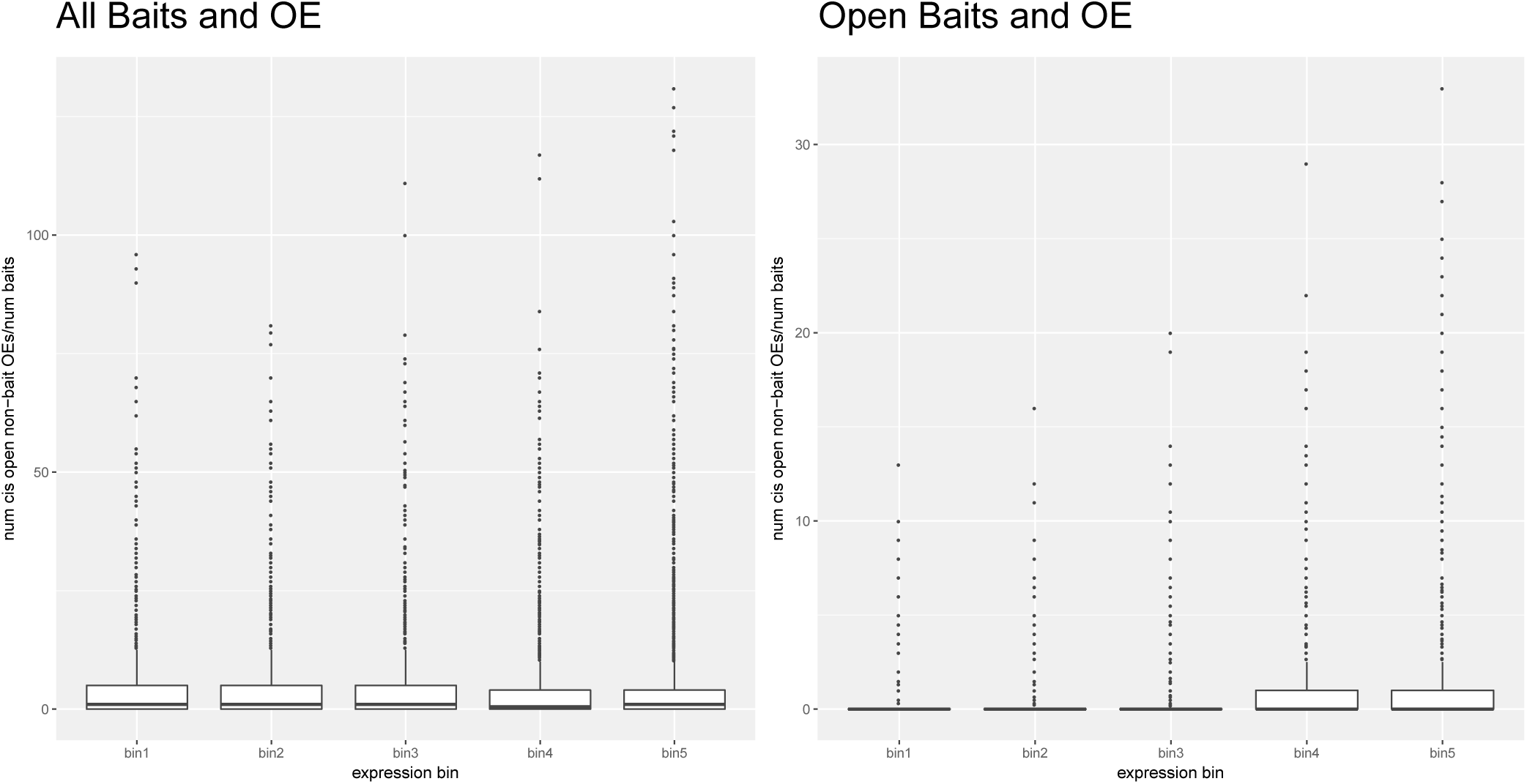

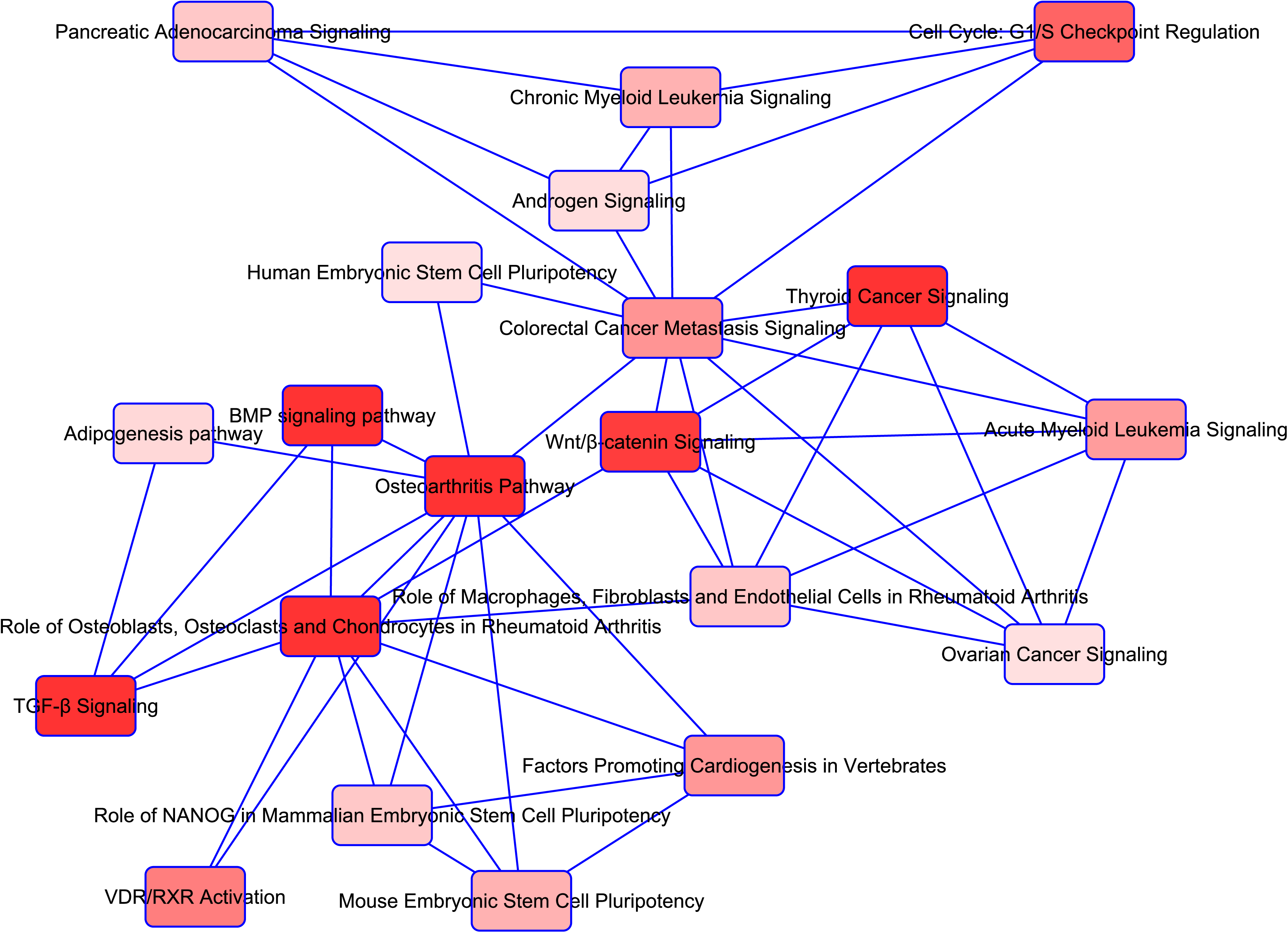

## ACKNOWLEDGEMENTS

This research was funded by the Children’s Hospital of Philadelphia. Some of the materials employed in this work were provided by the Texas A&M Health Science Center College of Medicine Institute for Regenerative Medicine at Scott & White through a grant from ORIP of the NIH, Grant # P40OD011050.

## References

1. Clark, E.M., Ness, A.R., Bishop, N.J. & Tobias, J.H. Association between bone mass and fractures in children: a prospective cohort study. J Bone Miner Res 21, 1489–95 (2006).

2. Goulding, A., Jones, I.E., Taylor, R.W., Williams, S.M. & Manning, P.J. Bone mineral density and body composition in boys with distal forearm fractures: a dual-energy x-ray absorptiometry study. J Pediatr 139, 509–15 (2001).

3. Kalkwarf, H.J., Laor, T. & Bean, J.A. Fracture risk in children with a forearm injury is associated with volumetric bone density and cortical area (by peripheral QCT) and areal bone density (by DXA). Osteoporos Int 22, 607–16 (2011).

4. Cauley, J.A. Public health impact of osteoporosis. J Gerontol A Biol Sci Med Sci 68, 1243–51 (2013).

5. Burge, R. et al. Incidence and economic burden of osteoporosis-related fractures in the United States, 2005–2025. J Bone Miner Res 22, 465–75 (2007).

6. Seeman, E. et al. Reduced bone mass in daughters of women with osteoporosis. N Engl J Med 320, 554–8 (1989).

7. Soroko, S.B., Barrett-Connor, E., Edelstein, S.L. & Kritz-Silverstein, D. Family history of osteoporosis and bone mineral density at the axial skeleton: the Rancho Bernardo Study. J Bone Miner Res 9, 761–9 (1994).

8. Duren, D.L. et al. Quantitative genetics of cortical bone mass in healthy 10-year-old children from the Fels Longitudinal Study. Bone 40, 464–70 (2007).

9. Heaney, R.P. et al. Peak bone mass. Osteoporos Int 11, 985–1009 (2000).

10. Krall, E.A. & Dawson-Hughes, B. Heritable and life-style determinants of bone mineral density. J Bone Miner Res 8, 1–9 (1993).

11. Bachrach, L.K., Hastie, T., Wang, M.C., Narasimhan, B. & Marcus, R. Bone mineral acquisition in healthy Asian, Hispanic, black, and Caucasian youth: a longitudinal study. Journal of Clinical Endocrinology and Metabolism 84, 4702–12 (1999).

12. Zemel, B.S. et al. Revised reference curves for bone mineral content and areal bone mineral density according to age and sex for black and non-black children: results of the bone mineral density in childhood study. J Clin Endocrinol Metab 96, 3160–9 (2011).

13. Grant, S.F. et al. Reduced bone density and osteoporosis associated with a polymorphic Sp1 binding site in the collagen type I alpha 1 gene. Nat Genet 14, 203–5 (1996).

14. Morrison, N.A. et al. Prediction of bone density from vitamin D receptor alleles. Nature 367, 284–7 (1994).

15. Styrkarsdottir, U. et al. Linkage of osteoporosis to chromosome 20p12 and association to BMP2. PLoS Biol 1, e69 (2003).

16. Little, R.D. et al. A mutation in the LDL receptor-related protein 5 gene results in the autosomal dominant high-bone-mass trait. Am J Hum Genet 70, 11–9 (2002).

17. Styrkarsdottir, U. et al. Multiple genetic loci for bone mineral density and fractures. N Engl J Med 358, 2355–65 (2008).

18. Richards, J.B. et al. Bone mineral density, osteoporosis, and osteoporotic fractures: a genome-wide association study. Lancet 371, 1505–12 (2008).

19. Styrkarsdottir, U. et al. New sequence variants associated with bone mineral density. Nat Genet 41, 15–7 (2009).

20. Kiel, D.P. et al. Genome-wide association with bone mass and geometry in the Framingham Heart Study. BMC Medical Genetics 8, S14 (2007).

21. Rivadeneira, F. et al. Twenty bone-mineral-density loci identified by large-scale meta-analysis of genome-wide association studies. Nat Genet 41, 1199–206 (2009).

22. Estrada, K. et al. Genome-wide meta-analysis identifies 56 bone mineral density loci and reveals 14 loci associated with risk of fracture. Nat Genet 44, 491–501 (2012).

23. Chesi, A. et al. A Genomewide Association Study Identifies Two Sex-Specific Loci, at SPTB and IZUMO3, Influencing Pediatric Bone Mineral Density at Multiple Skeletal Sites. J Bone Miner Res 32, 1274–1281 (2017).

24. Chesi, A. et al. A trans-ethnic genome-wide association study identifies gender-specific loci influencing pediatric aBMD and BMC at the distal radius. Hum Mol Genet 24, 5053–9 (2015).

25. Medina-Gomez, C. et al. BMD Loci Contribute to Ethnic and Developmental Differences in Skeletal Fragility across Populations: Assessment of Evolutionary Selection Pressures. Mol Biol Evol 32, 2961–72 (2015).

26. Mitchell, J.A. et al. Genetics of Bone Mass in Childhood and Adolescence: Effects of Sex and Maturation Interactions. J Bone Miner Res 30, 1676–83 (2015).

27. Kemp, J.P. et al. Phenotypic dissection of bone mineral density reveals skeletal site specificity and facilitates the identification of novel loci in the genetic regulation of bone mass attainment. PLoS Genet 10, e1004423 (2014).

28. Mitchell, J.A. et al. Rare EN1 Variants and Pediatric Bone Mass. J Bone Miner Res 31, 1513–7 (2016).

29. Zheng, H.F. et al. Whole-genome sequencing identifies EN1 as a determinant of bone density and fracture. Nature (2015).

30. Smemo, S. et al. Obesity-associated variants within FTO form long-range functional connections with IRX3. Nature 507, 371–5 (2014).

31. Claussnitzer, M. et al. FTO Obesity Variant Circuitry and Adipocyte Browning in Humans. N Engl J Med 373, 895–907 (2015).

32. Rosen, C.J. & Ingelfinger, J.R. Unraveling the Function of FTO Variants. N Engl J Med 373, 964–5 (2015).

33. Xia, Q. et al. The type 2 diabetes presumed causal variant within TCF7L2 resides in an element that controls the expression of ACSL5. Diabetologia 59, 2360–2368 (2016).

34. Grant, S.F. et al. Variant of transcription factor 7-like 2 (TCF7L2) gene confers risk of type 2 diabetes. Nat Genet 38, 320–3 (2006).

35. Replication, D.I.G. et al. Genome-wide trans-ancestry meta-analysis provides insight into the genetic architecture of type 2 diabetes susceptibility. Nat Genet 46, 234–44 (2014).

36. Consortium, G.T. et al. Genetic effects on gene expression across human tissues. Nature 550, 204–213 (2017).

37. Zheng, H.F. et al. Whole-genome sequencing identifies EN1 as a determinant of bone density and fracture. Nature 526, 112–7 (2015).

38. Medina-Gomez, C. et al. Bivariate genome-wide association meta-analysis of pediatric musculoskeletal traits reveals pleiotropic effects at the SREBF1/TOM1L2 locus. Nat Commun 8, 121 (2017).

39. Javierre, B.M. et al. Lineage-Specific Genome Architecture Links Enhancers and Non-coding Disease Variants to Target Gene Promoters. Cell 167, 1369–1384 e19 (2016).

40. Hughes, J.R. et al. Analysis of hundreds of cis-regulatory landscapes at high resolution in a single, high-throughput experiment. Nat Genet 46, 205–12 (2014).

41. Zhu, F., Friedman, M.S., Luo, W., Woolf, P. & Hankenson, K.D. The transcription factor osterix (SP7) regulates BMP6-induced human osteoblast differentiation. J Cell Physiol 227, 2677–85 (2012).

42. Cairns, J. et al. CHiCAGO: robust detection of DNA looping interactions in Capture Hi-C data. Genome Biol 17, 127 (2016).

43. Consortium, E.P. An integrated encyclopedia of DNA elements in the human genome. Nature 489, 57–74 (2012).

44. Tiaden, A.N. et al. Human serine protease HTRA1 positively regulates osteogenesis of human bone marrow-derived mesenchymal stem cells and mineralization of differentiating bone-forming cells through the modulation of extracellular matrix protein. Stem Cells 30, 2271–82 (2012).

45. Jin, C. et al. Inhibition of lncRNA MIR31HG Promotes Osteogenic Differentiation of Human Adipose-Derived Stem Cells. Stem Cells 34, 2707–2720 (2016).

46. Baxter, J.S. et al. Capture Hi-C identifies putative target genes at 33 breast cancer risk loci. Nat Commun 9, 1028 (2018).

47. Lieberman-Aiden, E. et al. Comprehensive mapping of long-range interactions reveals folding principles of the human genome. Science 326, 289–93 (2009).

48. Mifsud, B. et al. Mapping long-range promoter contacts in human cells with high-resolution capture Hi-C. Nat Genet 47, 598–606 (2015).

49. Mumbach, M.R. et al. Enhancer connectome in primary human cells identifies target genes of disease-associated DNA elements. Nat Genet 49, 1602–1612 (2017).

50. Nabbi, A. et al. ING3 protein expression profiling in normal human tissues suggest its role in cellular growth and self-renewal. Eur J Cell Biol 94, 214–22 (2015).

51. Nagashima, M. et al. A novel PHD-finger motif protein, p47ING3, modulates p53-mediated transcription, cell cycle control, and apoptosis. Oncogene 22, 343–50 (2003).

52. Ganss, B. & Hoffmann, W. Calcium-induced conformational transition of trout ependymins monitored by tryptophan fluorescence. Open Biochem J 3, 14–7 (2009).

53. Moverare-Skrtic, S. et al. Osteoblast-derived WNT16 represses osteoclastogenesis and prevents cortical bone fragility fractures. Nat Med 20, 1279–88 (2014).

54. Haraguchi, R. et al. sFRP4-dependent Wnt signal modulation is critical for bone remodeling during postnatal development and age-related bone loss. Sci Rep 6, 25198 (2016).

55. Kiper, P.O.S. et al. Cortical-Bone Fragility-Insights from sFRP4 Deficiency in Pyle’s Disease. N Engl J Med 374, 2553–2562 (2016).

56. Medina-Gomez, C. et al. Life-Course Genome-wide Association Study Meta-analysis of Total Body BMD and Assessment of Age-Specific Effects. Am J Hum Genet 102, 88–102 (2018).

57. Mitchell, J.A. et al. Multidimensional Bone Density Phenotyping Reveals New Insights Into Genetic Regulation of the Pediatric Skeleton. J Bone Miner Res 33, 812–821 (2018).

58. Luo, W., Friedman, M.S., Hankenson, K.D. & Woolf, P.J. Time series gene expression profiling and temporal regulatory pathway analysis of BMP6 induced osteoblast differentiation and mineralization. BMC Syst Biol 5, 82 (2011).

59. Wingett, S. et al. HiCUP: pipeline for mapping and processing Hi-C data. F1000Res 4, 1310 (2015).

60. Dobin, A. et al. STAR: ultrafast universal RNA-seq aligner. Bioinformatics 29, 15–21 (2013).

